# Divergent hepaciviruses, chuvirus and deltaviruses in Australian marsupial carnivores (Dasyurids) identified through transcriptome mining

**DOI:** 10.1101/2023.06.27.546737

**Authors:** Erin Harvey, Jonathon C.O. Mifsud, Edward C. Holmes, Jackie E. Mahar

## Abstract

Although Australian marsupials are characterised by unique biology and geographic isolation, little is known about the viruses present in these iconic wildlife species. The Dasyuromorphia are an order of marsupial carnivores found only in Australia that include both the extinct Tasmanian tiger (Thylacine) and the highly threatened Tasmanian devil. Several other members of the order are similarly under threat of extinction due to habitat loss, hunting, disease, and competition and predation by introduced species such as feral cats. We utilised publicly available RNA-seq data from the NCBI Sequence Read Archive (SRA) database to document the viral diversity within four Dasyuromorphia species. Accordingly, we identified 15 novel virus species from five DNA virus families (*Adenoviridae*, *Anelloviridae*, *Herpesviridae*, *Papillomaviridae* and *Polyomaviridae*) and three RNA virus taxa: the order *Jingchuvirales,* the genus *Hepacivirus*, and the delta-like virus group. Of particular note was the identification of a marsupial specific clade of delta-like viruses that may indicate an association of deltaviruses and with marsupial species dating back to their origin some 160 million years ago. In addition, we identified a highly divergent hepacivirus in a numbat liver transcriptome that falls outside of the larger mammalian clade, as well as the first detection of the *Jingchuvirales* in a mammalian host – a chu-like virus in Tasmanian devils – thereby expanding the host range beyond invertebrates and ectothermic vertebrates. As many of these Dasyuromorphia species are currently being used in translocation efforts to reseed populations across Australia, understanding their virome is of key importance to prevent the spread of viruses to naive populations.

## Introduction

Australian wildlife have evolved in isolation for approximately 45 million years, resulting in a unique mammalian fauna of which 87% are endemic (Chapman 2009). This includes carnivorous marsupials of the order Dasyuromorphia (Kealy and Beck 2017). Members of this order have experienced extinction as a direct result of human activity, with the Tasmanian tiger *(Thylacinus cynocephalus*) an iconic symbol of human-mediated mammalian extinction (Feigin et al. 2022). Indeed, Australia is experiencing one of the highest rates of mammalian extinction globally due to changing land-use, disease, and competition and predation from introduced species (Woinarski et al. 2011). Despite these threats, we know little about the viruses that infect these unique and at-risk species and how these viruses might impact population health.

Marsupials are an infraclass of mammals, characterised by their distinctive pouch in which they carry live young. The Dasyuromorphia represent all carnivorous marsupials in Australia apart from the omnivorous bandicoots (Peramelemorphia) (Zemann et al. 2013), and are found only on the mainland of Australia and its surrounding islands, such as the Australian island state of Tasmania, and Papua New Guinea (Kealy and Beck 2017). The largest marsupial carnivore was the extinct thylacine, followed by the now endangered Tasmanian devil (*Sarcophilus harrisii*) that has replaced the tiger as the apex predator in Tasmania. Through a combination of hunting, habitat loss, competition for resources with introduced species and potentially the introduction of exotic pathogens, the last known thylacine died in 1936 and the species was declared officially extinct in 1986 (Paddle 2002). The Tasmanian devil is also threatened by a contagious cancer that has contributed to an average population decline of 77% in the past 20 years (Lazenby et al. 2018).

Beyond these two iconic species, lesser-known marsupial carnivores are also at risk of extinction. The numbat (*Myrmecobius fasciatus*) is the last remaining member of the *Myrmecobiidae*, the only diurnal marsupial, and the only marsupial that feeds exclusively on termites (Zemann et al. 2013). Due to habitat loss and predation by introduced species such as cats and foxes, the numbat only persists in two natural locations in southwestern Australia and there are estimated to be less than 1000 numbats remaining in the wild (Peel et al. 2022). While other species that had been believed to be of little concern such as the fat-tailed dunnart (*Sminthopsis crassicaudata*) have recently been listed as threatened by the state of Victoria due to population decline (Scicluna et al. 2021). To combat these dramatic declines in populations of marsupial carnivores across Australia, programs have been employed to relocate individuals from thriving populations in protected locations, such as the quoll populations in Tasmania, as well as from captive breeding populations (Portas et al. 2020). Similarly, the yellow-footed antechinus (*Antechinus flavipes*) has been proposed as a candidate species for translocation as part of a rewilding project of an area near Canberra, Australia (Manning 2021). These programs have experienced varied success, and an often overlooked risk with the translocation of wildlife is the introduction of pathogens into naive populations already struggling under the burden of anthropogenic activities and introduced species (French et al. 2022).

We know little about the viruses circulating within dasyuromorphs. To date, only a single virome project has been performed on these animals – in this case the faecal virome of Tasmanian devils (Chong et al. 2019). There have been a small number of studies of marsupial carnivores experiencing overt signs of disease, including the characterisation of Dasyurid herpesvirus 1 in *Antechinus* (Barker et al. 1981), and a papilloma-polyomavirus hybrid has been identified in bandicoots (Woolford et al. 2007). There have also been a small number of serological and bioinformatic screening studies targeting or including marsupial carnivores. For example, the serological screening of marsupial tissues for herpesviruses led to the identification of a novel herpes virus in a Tasmanian devil (Stalder et al. 2015), while an analysis of available transcriptomic data and reference host genomes characterised the endogenous viral elements (EVEs) of marsupial carnivores (Harding et al. 2021).

Herein, we present the first systematic attempt to identify viruses in marsupial carnivores. Because of the inherent challenges in acquiring samples from dasyuromorphs that are generally protected species, we instead mined the available transcriptomes present on the NCBI Sequence Read Archive (SRA), following by genomic and phylogenetic analysis.

## 2. Methods

### 2.1. Identification of virus contigs in transcriptome data

A custom virus detection pipeline was used to screen all available Dasyuromorphia (NCBI taxonomic identifier taxid 2759) SRA RNA-seq data sets, excluding faecal samples. Raw FASTQ files for all libraries were downloaded using Kingfisher (https://github.com/wwood/kingfisher-download). Sequencing reads first underwent quality trimming and adapter removal using Trimmomatic (v0.38) with parameters SLIDINGWINDOW:4:5, LEADING:5, TRAILING:5, and MINLEN:25, prior to assembly (Bolger et al. 2014).

*De novo* assembly was conducted using MEGAHIT with default parameters (v1.2.9) (D. Li et al. 2015). The assembled contigs were then compared to the RdRp-scan RNA-dependent RNA polymerase (RdRp) core protein sequence database (v0.90) (Charon et al. 2022) and the protein version of the Reference Viral Databases (v23.0) (Bigot et al. 2019; Goodacre et al. 2018) using Diamond BlastX (v2.0.9) with an e-value cut-off of 1 × 10−5 (Buchfink et al. 2021). To remove potential false positives, contigs with hits to virus sequences were used as a query against the NCBI nucleotide database (as of May 2022) using Blastn, and all contigs with sequence similarity to non-virus nucleotide sequences were removed from the query set (Camacho et al. 2009). The remaining contigs were then aligned against the NCBI non-redundant protein database (as of March 2022) using Diamond BlastX and contigs with hits to virus proteins were further examined. The *de novo* assembler and Find ORFs tool, available in Geneious (Kearse et al. 2012), were used to further assemble genomes where necessary and to identify ORFs in potential virus sequences, respectively. EVEs were identified by referencing Harding et al. 2021 and removed manually. NCBI web blast (https://www.ncbi.nlm.nih.gov/BLAST) was then used to check for false positives, disrupted ORFs, and to manually assess alignment to virus motifs.

### 2.3. Virus abundance

The abundance of virus contigs was measured using the RNA-seq by Expectation Maximisation (RSEM) (v1.3.0) program (B. Li and Dewey 2011). Expected count of viral contigs was used to calculate viral abundance as a percentage of total reads. All calculations and graphing were performed using R.

### 2.4. Phylogenetic analysis

Sequences representing novel virus species, defined as those with <95% nucleotide sequence similarity to their closest relative, were assigned to a taxonomic family based on their similarity to previously characterised virus species. For each family, a reference data set was downloaded from NCBI Virus (Hatcher et al. 2017) and the completeness of this data set was assessed by comparing it to the ICTV-recognised species for each family. Unclassified species related to the viruses discovered here were added using a web Blast search using the novel virus sequence as the query. Amino acid alignments were performed for all families except *Kolmioviridae* (Deltavirus) due to the high level of divergence within the reference data set. Multiple sequence alignments were produced using MAFFT (v 7.402) (Katoh and Standley 2013) with local pair alignment for amino acid sequences and global pair alignment for the deltavirus nucleotide sequences. All alignments were then trimmed to remove ambiguous regions using trimAl (v1.4.1) (Capella-Gutierrez et al. 2009) with a gap threshold of 0.8 and a similarity threshold of 0.005, and then manually assessed using AliView (Larsson 2014). Phylogenetic trees were estimated using IQ-TREE 2 (v2.2.2) (Minh et al. 2020) with the appropriate substitution model selected using ModelFinder (Kalyaanamoorthy et al. 2017). Branch support was assessed with 1000 bootstrap replicates using ultrafast bootstrapping (Hoang et al. 2017).

### 2.5. Mapping sequences to *Herpes virus*

As only one reference sequence for a single gene (DNA dependent DNA polymerase) was available for the species of *Antechinus* herpesvirus previously identified, Bowtie2 (v2.2.5) (Langmead and Salzberg 2012) was used to align the non-host reads to the Dasyurid herpesvirus 1 DNA-DNA polymerase reference sequence (MF576269.1). The resulting alignment was viewed in Geneious (Kearse et al. 2012) and used to assess the percentage sequence similarity between the novel sequence and the available reference.

### 2.6. Library composition assessment

To assess the taxonomic composition of each library, contigs were aligned to a custom NCBI nucleotide database without environmental and artificial sequences (https://researchdata.edu.au/indexed-reference-databases-kma-ccmetagen/1371207) using the KMA aligner (v1.3.9a) and the CCMetagen program (v1.1.3) (Clausen et al., 2018; Marcelino et al., 2020). The abundance of each taxonomic group was determined by counting the number of nucleotides that matched the reference sequence, with an additional correction for template length using the default parameter in KMA. For data visualization, CCMetagen was used to generate Krona graphs, which were subsequently edited in Adobe Illustrator (https://www.adobe.com).

## 3. Results

### 3.1. Screening Dasyuromorphia for viruses

As of March 2023, the NCBI SRA database contained 446 RNA-seq libraries from *Dasyuromorphia* (Supp. Table 1), comprising seven species: *Sminthopsis crassicaudata* (fat-tailed dunnart), *Antechinus flavipes* (yellow-footed antechinus), *Antechinus stuartii* (brown antechinus), *Sarcophilus harrisii* (Tasmanian devil), *Thylacinus cynocephalus* (Tasmanian tiger), *Myrmecobius fasciatus* (numbat) and *Pseudantechinus macdonnellensis* (fat-tailed false antechinus).

Virus sequence similarity screening of the available libraries resulted in potential positive hits in 206 of the screened libraries, including Tasmanian devil, yellow-footed antechinus, numbut, and fat-tailed dunnart. Following secondary assembly and removal of contigs with hits to endogenous viruses, 22 partial or full virus genomes were identified in 43 libraries including four host species (Figure 1). These viruses were assigned taxonomically to five DNA virus families: *Adenoviridae*, *Anelloviridae*, *Herpesviridae*, *Papillomaviridae* and *Polyomaviridae*, and three RNA virus taxa: the order *Jingchuvirales,* the genus *Hepacivirus*, and the Delta-like virus group (Table 1). Of these, two were partial sequences of novel herpesviruses in Tasmanian devil and the fat-tailed dunnart that were too short for phylogenetic confirmation and thus excluded from further study. There was also a fragment of the ORF2 of a novel *Anellovirus* present in a Tasmanian devil library, but as the ORF1 fragment which is used for phylogenetic classification of these viruses was absent, and ORF2 is rarely published for this virus species, it was similarly excluded.

**Figure 1.**
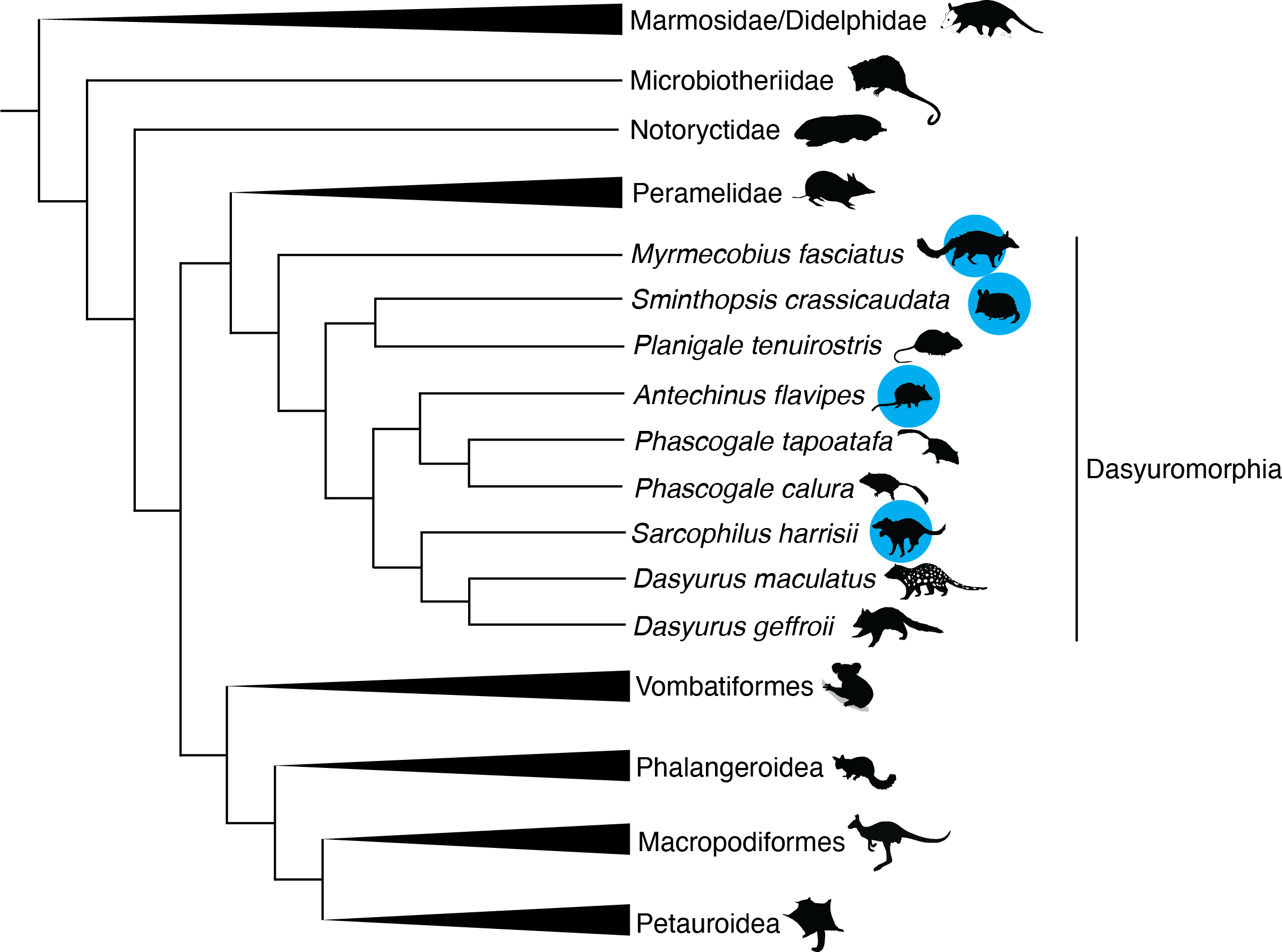
Cladogram depicting the phylogenetic relationships of marsupials. Adapted from Duchene et al 2017 (Duchêne et al. 2017). Species in which viruses were identified in this study are highlighted with blue circles. Animal silhouettes are sourced from PhyloPic (https://www.phylopic.org/) produced by Sarah Werning, Gabriela Palomo-Munoz. Creative commons licence found at https://creativecommons.org/licenses/by/3.0/.

**Table 1.** Viruses identified in this study including the details of the SRA libraries in which they were identified.

| Virus name | Closest blast hit (nr) | % similarity to top blast hit (aa) | Virus Taxonomy | Host | Tissue | BioProject | Library ID |
| --- | --- | --- | --- | --- | --- | --- | --- |
| Fat-tailed dunnart deliavirus | Rodent deltavirus (QJD13562.1) | 71 | *Deltavirus* | *Sminthopsis crassicaudata* | Adult eye | PRJNA554238 | SRR9673767 |
| Tasmanian devil deltavirus | Rodent deltavirus (QJD13562.1) | 71 | *Deltavirus* | *Sarcophilus harrisii* | Tasmanian devil facial tumour | PRJNA693818 | SRR13765777 |
| Tasmanian devil chu-like virus | Lishi spider virus 1 (AJG39051.1) | 24 | *Jingchuvirales* | *Sarcophilus harrisii* | Tasmanian Devil facial tumour (DFTD1) cell lines | PRJNA422607 | SRR6380970 |
| Antechinus hepacivirus | Northern treeshrew hepacivirus (CAI5760841.1) | 48 | *Hepacivirus* | *Antechinus flavipes* | Spleen, kidney, liver, stomach, cerebrum | PRJNA565840 | SRR10127601 SRR10127603<br>SRR11306583 SRR11306585<br>SRR11306588 SRR11306615<br>SRR11306617 SRR11306621<br>SRR11306628 SRR11306634<br>SRR11306639 SRR11306641<br>SRR11306643 SRR11306667<br>SRR11306686 |
| Numbat hepacivirus | Duck hepacivirus (QKT21547.1) | 29 | *Hepacivirus* | *Myrmecobius fasciatus* | Lung, Liver | PRJNA786364 | SRR17244188 |
| Dasyurid herpesvirus 4 | Ursid gammaherpesvirus 2 (AZP55492.1) | 37 | *Herpesviridae* | *Antechinus flavipes* | male prostate | PRJNA565840 | SRR11306664 |
| Dasyurid herpesvirus 5 | Ursid gammaherpesvirus 2 (AZP55492.1) | 38 | *Herpesviridae* | *Antechinus flavipes* | Male stomach | PRJNA565840 | SRR11306624 |
| Tasmanian devil polyomavirus 3 | Goose hemorrhagic polyomavirus (UJT42146.1) | 43 | *Polyomaviridae* | *Sarcophilus harrisii* | Lip tissue | PRJNA693818 | SRR13765816 SRR13765794 |
| Antechinus Anellovirus | Torque teno virus (QBA84075.1) | 28 | *Anelloviridae* | *Antechinus flavipes* | spleen | PRJNA565840 | SRR11306585 SRR11306684 |
| Antechinus adenovirus | Bat mastadenovirus WIV13 (YP_009272931.1) | 69 | *Adenoviridae* | *Antechinus flavipes* | Kidney, prostate, spleen | PRJNA565840 | SRR11306638 SRR11306640<br>SRR11306641 SRR11306643<br>SRR11306644 SRR11306646<br>SRR11306647 SRR11306648<br>SRR11306664 SRR11306685 |
| Tasmanian devil papillomavirus 3 | Human papillomavirus type 229 (UQF78855.1) | 45 | *Papillomaviridae* | *Sarcophilus harrisii* | Lip tissue | PRJNA693818 | SRR13765825 |
| Tasmanian devil papillomavirus 4 | Capra hircus papillomavirus type 2 (QIH12244.1) | 40 | *Papillomaviridae* | *Sarcophilus harrisii* | Lip tissue | PRJNA693818 | SRR13765801 |
| Tasmanian devil papillomavirus 5 | Human papillomavirus (QAB13979.1) | 43 | *Papillomaviridae* | *Sarcophilus harrisii* | Lip tissue | PRJNA693818 | SRR13765803 SRR13765816 |
| Tasmanian devil papillomavirus 6 | Manis javanica papillomavirus 1 (DAZ92261.1) | 47 | *Papillomaviridae* | *Sarcophilus harrisii* | Lip tissue | PRJNA693818 | SRR13765808 |
| Tasmanian devil papillomavirus 7 | Bos taurus papillomavirus 20 (YP_009272609.1) | 40 | *Papillomaviridae* | *Sarcophilus harrisii* | Lip tissue | PRJNA693818 | SRR13765811 |

Within the virus positive data set, further analysis revealed that four *Antechinus* libraries from the same study (SRR11306636, SRR11306642, SRR11306648 and SRR11306672) appeared to contain viruses identified as likely contaminants. Specifically, the human associated viruses Gammapapillomavirus 19 and Betapapillomavirus 1 were identified with over 99% nucleotide sequence similarity, and human reads were found in three of these libraries (Supp. Fig. 1.). These viruses were excluded from further analysis. No other libraries contained contaminating reads (Supp. Fig. 1).

### 3.2. Marsupial delta-like viruses

Of particular note, we identified the first marsupial associated delta-like viruses. These were detected in a fat-tailed dunnart and a Tasmanian devil, provisionally named Fat-tailed dunnart deltavirus and Tasmanian devil deltavirus, respectively. These viruses shared 71% amino acid sequence similarity with their nearest relative, Rodent deltavirus, but were also distinct from each other with 84% nucleotide similarity across the entire assembled contigs. Notably, these viruses formed a distinct clade within the Deltavirus small delta antigen-like protein phylogeny (Figure 2), clustering most closely with viruses sampled from rodents.

**Figure 2.**
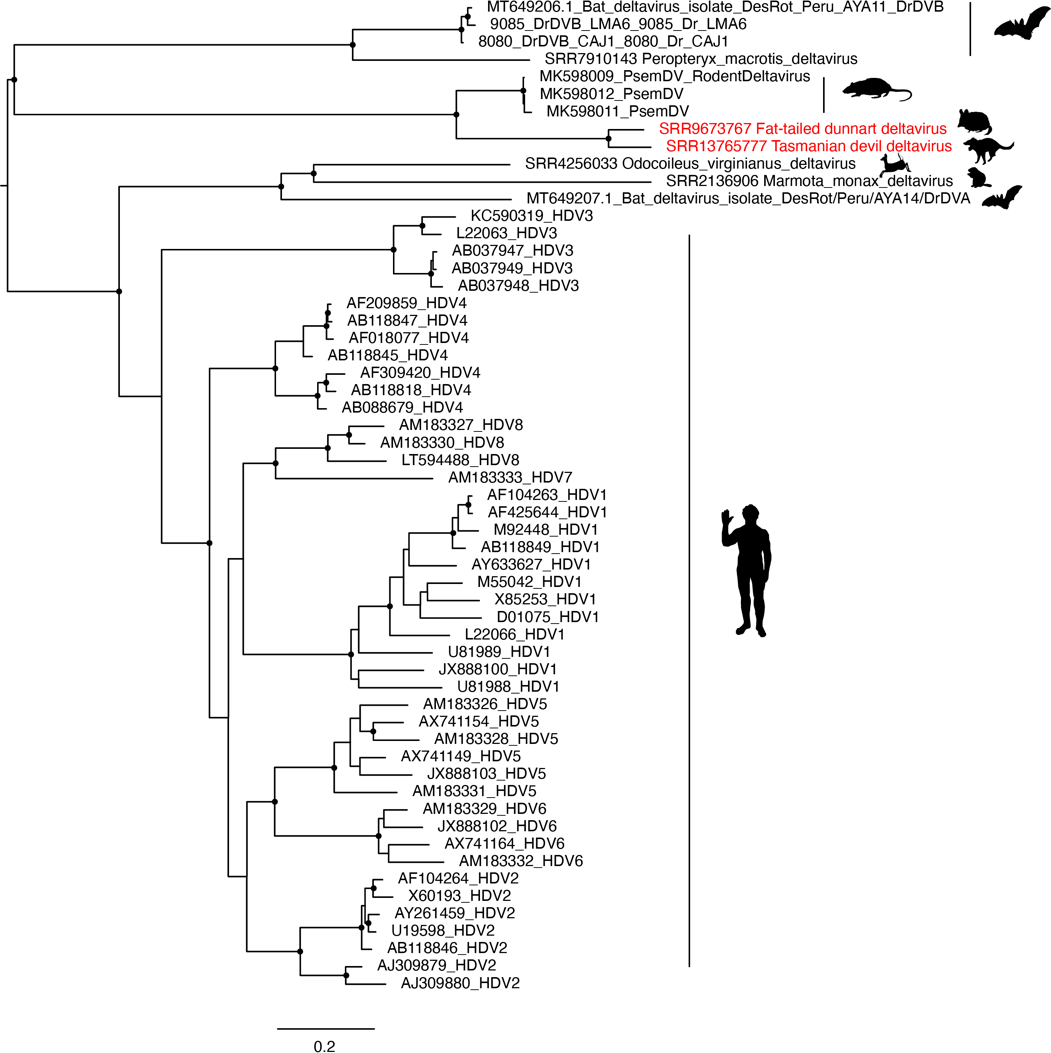
Phylogeny of mammalian deltavirus small delta antigen-like protein nucleotide sequence. The phylogeny was estimated using the GTR+F+I+ρ4 nucleotide substitution model. Sequences identified in this study are shown in red text and the library ID is indicated in the taxa name. Branch support >90% is indicated with a black dot at the node. The scale bar indicates the number of nucleotide substitutions per site and the tree is midpoint rooted for clarity. This alignment is based on that provided by Bergner et al. 2021. Animal silhouettes indicate the host species.

The fat-tailed dunnart deltavirus was identified in a library of eye tissue from a study of gene expression in mammalian embryos (Royall et al. 2019). The assembled contig was 643 nucleotides in length, thereby representing only a fragment of the expected 1.6kb viral genome, and was found at remarkably low abundance, with only 4.06e-7% of total reads (expected count: 272) aligning to the contig (Figure 3). Tasmanian devil deltavirus was identified in a sample of Tasmanian devil facial tumour disease taken from a female Tasmanian devil. The assembled contig was again fragmentary, at 741 nucleotides in length, and the abundance was also low with 3.98e-7% of reads (expected count: 27) aligning to the contig. In both cases, no other viruses were identified in the sequencing libraries.

**Figure 3.**
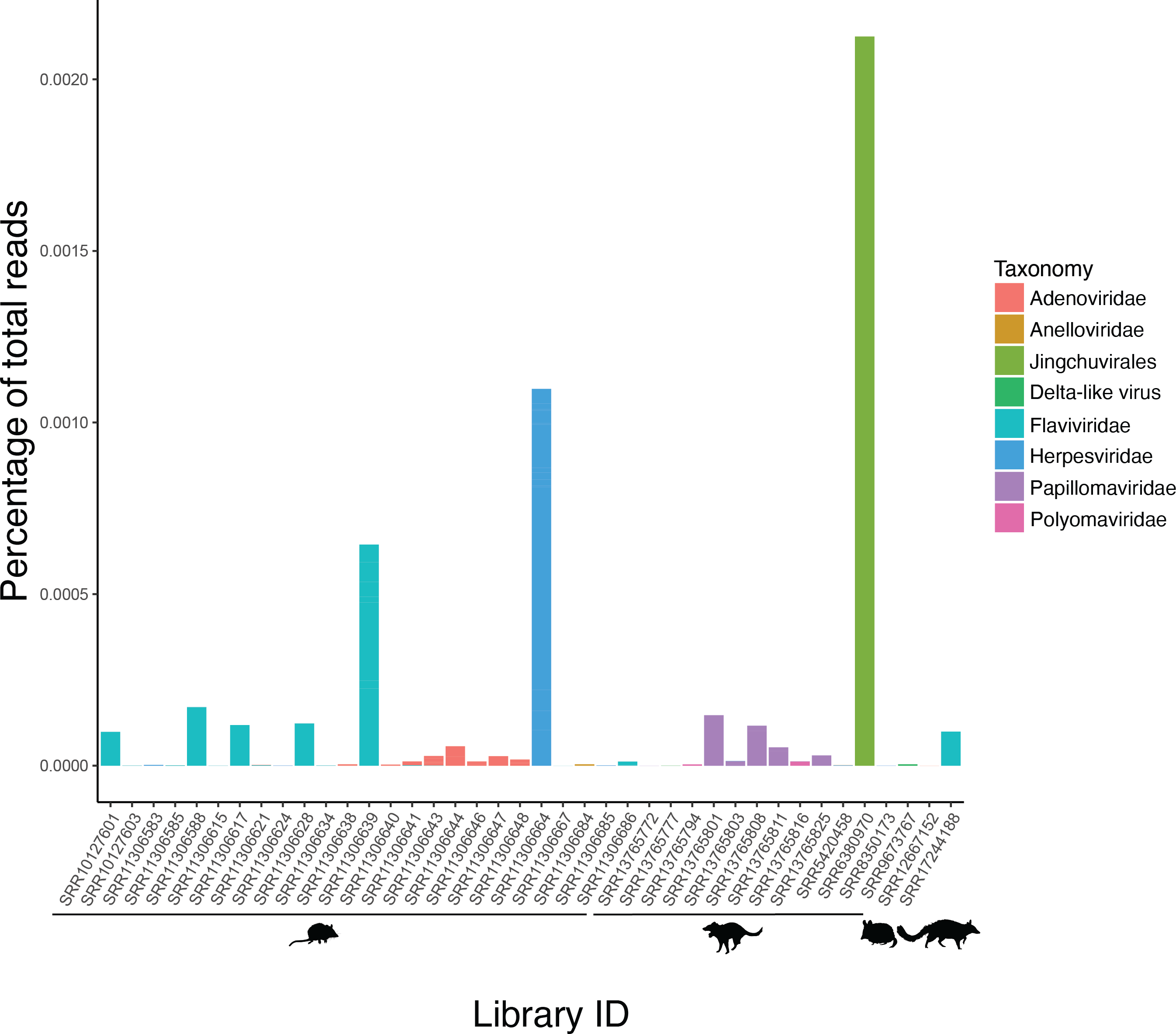
Virus abundance in marsupial carnivores as a percentage of total reads. The animal from which libraries were obtained is indicated by an animal silhouette below the library ID.

### 3.3. RNA viruses

We identified three species of novel RNA viruses in this study: two hepaciviruses (positive-sense RNA, family *Flaviviridae*), one in antechinus and one in a numbat, and a Chu-like virus (negative-sense RNA virus, order *Jingchuvirales*) here named Tasmanian devil chu-like virus. The chu-like virus was identified in a Devil facial tumour disease (DFTD) cell line that was cultured from primary tissue (Kozakiewicz et al. 2021), and was at the highest abundance of all viruses identified in this study, representing 0.2% of total reads (expected read count: 253,672) aligning to the viral genome (Figure 3). A phylogenetic analysis of the RdRp revealed that Tasmanian devil chu-like virus fell outside of the *Chuviridae* and was highly divergent from known viruses including the only known vertebrate clade of *Chuviridae*, the genus *Piscichuvirus*, although it did cluster within the order *Jingchuvirales* (Figure 4A). The high abundance in the cultured Tasmanian devil tissues and absence of any sequencing reads of a non-target source in this library (Bioproject SRR6380970) tentatively suggests that it does replicate in these cells and was not the result of contamination (Supp. Fig 1).

**Figure 4.**
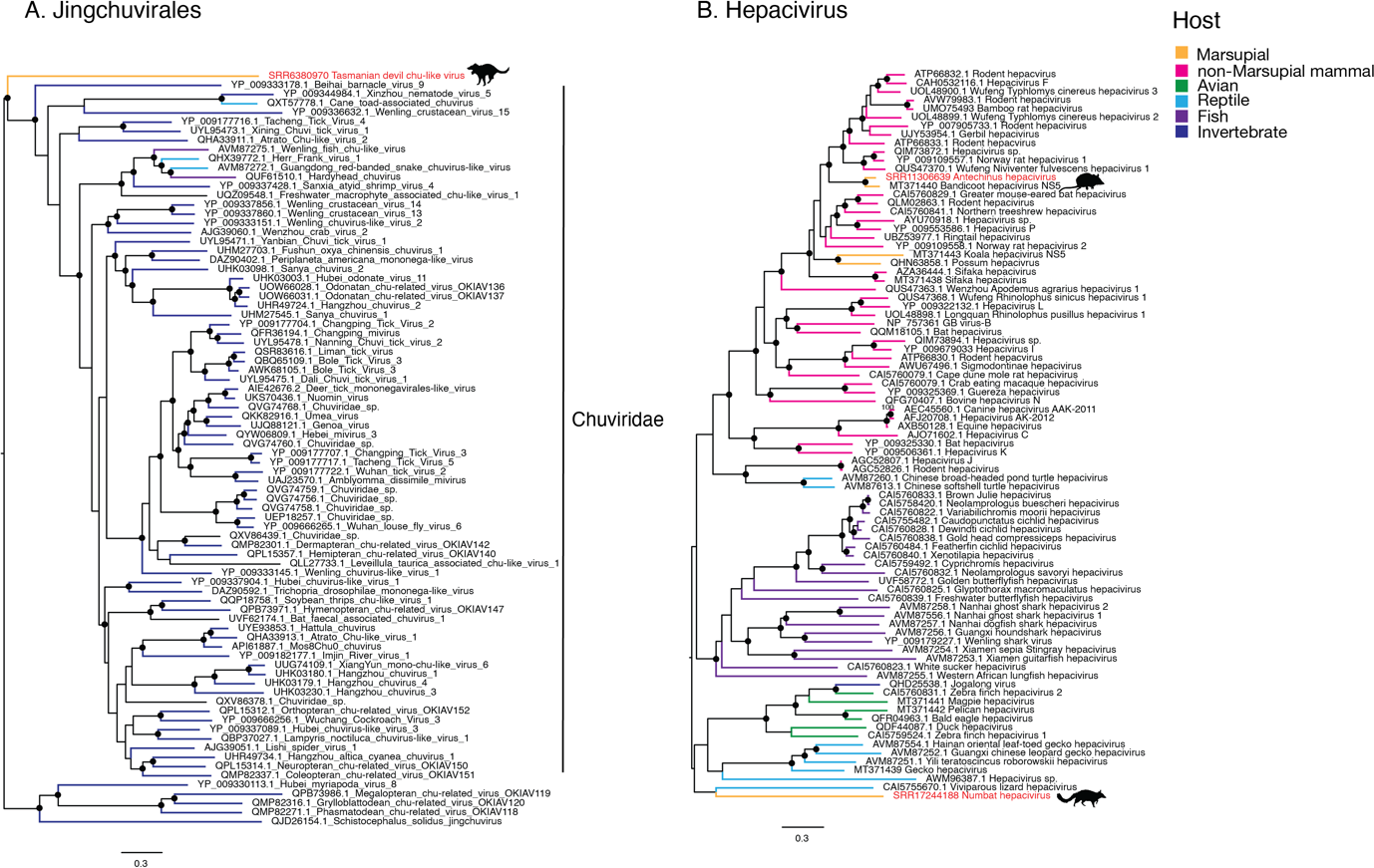
Phylogenetic trees of the RNA viruses identified in this study. (A) Phylogeny of the *Jingchuvirales* based on the RdRp amino acid sequence using the LG+F+I+ρ4 substitution model. (B) Phylogeny of the hepacivirus NS5 protein sequence estimated using the LG+F+I+ρ4 substitution model. Scale bars indicate the number of amino acid substitutions per site. Trees are midpoint rooted for clarity. Branch support values >90% are indicated with a black dot at the node. Viruses identified in this study are highlighted with red taxa labels. Animal silhouettes indicate the host species of the viruses identified here and tip branches are coloured according to broader host species.

Antechinus hepacivirus was detected in multiple libraries from a single BioProject (PRJNA565840) in which multiple tissues of 13 individual antechinus were sequenced. The assembled virus genome was 7209 nucleotides in length and was detected in five of the 13 individuals and in at least two tissue types from each positive individual, with libraries of liver tissue consistently having the highest abundance (1.18E-04 - 6.44E-04 percent of total reads). The virus was also identified in spleen, kidney, stomach, and cerebrum. Based on the NS5 (RdRp) protein, this virus was most closely related to a hepacivirus identified in an *Ixodes holocyclus* tick engorged with the blood of a long-nosed bandicoot (Harvey et al. 2019; Porter et al. 2020). In turn, these two viruses were related to rodent hepacivirus (Figure 4B).

Finally, a highly divergent hepacivirus was identified in a transcriptome of numbat liver tissue (Peel et al. 2022) with an abundance of 9.9e-5% of total reads. This virus, provisionally named Numbat hepacivirus, has 37% amino acid sequence similarity to the closest blast hit (Norway rat hepacivirus 2, YP_009325411.1) over the NS5 protein and 29% amino acid sequence similarity across the whole genome to the closest blast hit (Duck hepacivirus, QKT21547.1). Numbat hepacivirus clustered phylogenetically with a clade of avian and reptile associated hepaciviruses based on the NS5 protein (Figure 4B), but on a long branch characterised by low bootstrap support (63%) suggesting that its phylogenetic position is uncertain due to high sequence divergence.

### 3.4. DNA viruses

Although RNA-seq data was analysed here, our data set contained a surprising diversity of novel DNA virus transcripts. Herpesviruses were present in three species - antechinus, Tasmanian devil and fat-tailed dunnart - but only the antechinus libraries contained sufficient gene sequences for phylogenetic analysis and was therefore only these species were analysed further. A very small 55 nucleotide fragment of glycoprotein H of the herpesvirus identified in Tasmanian devils exhibited 100% nucleotide sequence similarity to Dasyurid herpesvirus 3 (Chong et al. 2019).

Herpesvirus sequences were identified in four Antechinus libraries – two from female spleen libraries containing two contigs each – although none were appropriate for phylogenetic analysis and did not overlap with the genes present in the other two libraries. These sequences were not investigated further. Within the remaining two libraries, one library from male stomach tissue (SRR11306624) contained two short contigs that included the glycoprotein B gene (as well as one hypothetical protein gene sequence) which could be utilised in phylogenetic analysis, here named Dasyurid herpesvirus 5. The other, a library of prostate tissue, contained 24 contigs with sequence similarity to gammaherpesvirus genes, also including the glycoprotein B sequence, here named Dasyurid herpesvirus 4. A comparison of the glycoprotein B sequence of these two viruses revealed that they shared only 25% amino acid sequence similarity. We used Bowtie2 to align reads from the herpesvirus positive libraries to the Dasyurid herpesvirus 1 reference sequence (MF576269.1), for which only the DNA-DNA polymerase is currently available. A single read from SRR11306664 (containing Dasyurid herpesvirus 4) aligned with 92% nucleotide sequence similarity to the Dasyurid herpesvirus 1 reference, while no reads aligned from the other three Antechinus libraries containing herpesvirus contigs. Based on phylogenetic analysis of glycoprotein B, Dasyurid herpesvirus 4 was most closely related to an elephant herpesvirus, but more broadly with vombatid herpesvirus and koala herpesvirus (Figure 5C). Dasyurid herpesvirus 5 fell as a sister lineage to this group, although the sequence is very short and characterised by low bootstrap support. Without the DNA-DNA polymerase, the most commonly sequenced Herpesvirus gene used for taxonomic demarcation, the taxonomic relationship of these viruses was uncertain.

**Figure 5.**
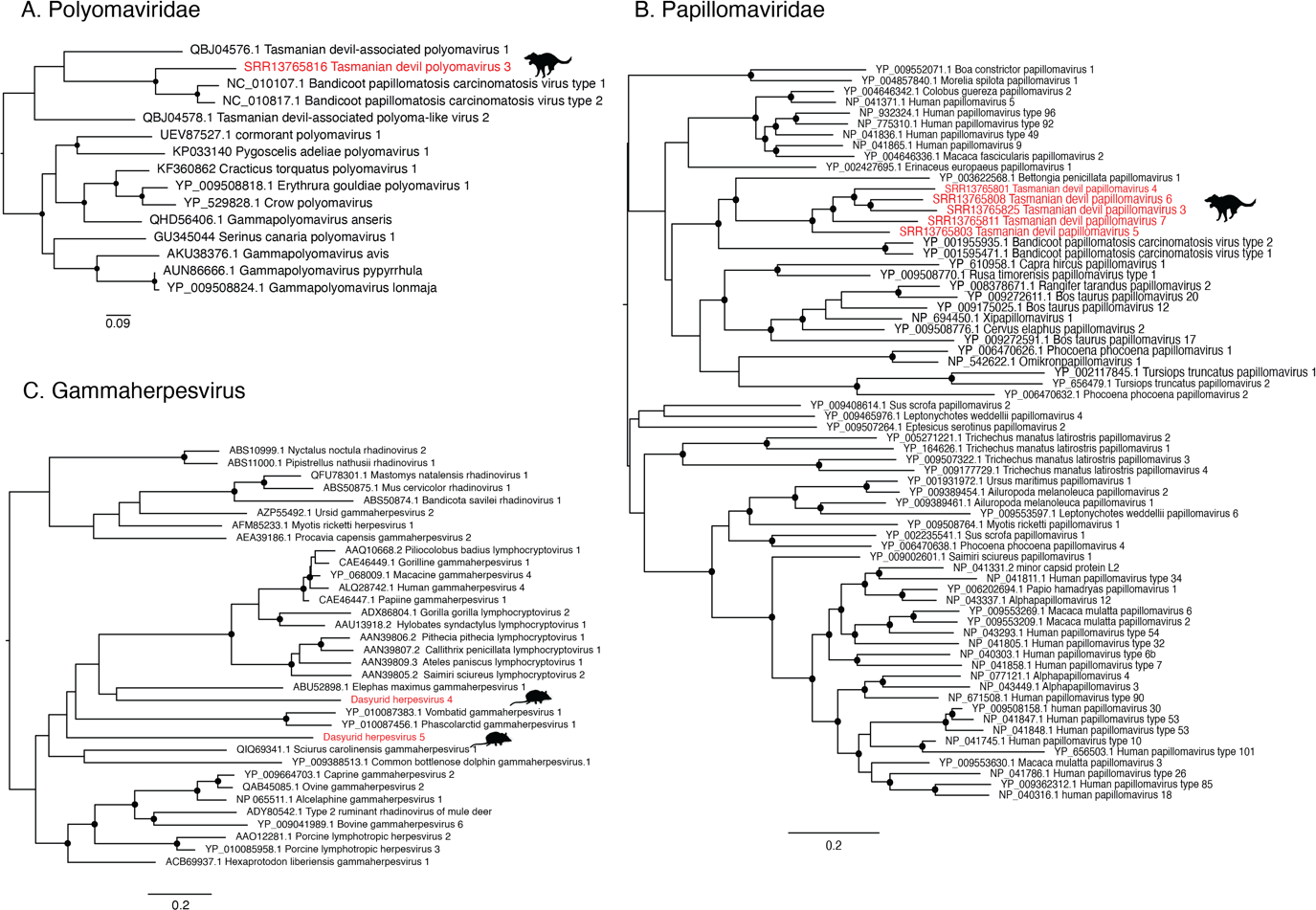
Phylogenies of *Papillomaviridae*, *Polyomaviridae* and *Gammaherpesviridae*. (A) Phylogeny of the *Polyomaviridae* large T antigen protein estimated using the LG+I+ρ4 substitution model. (B) Phylogeny of the *Papillomaviridae* L1 and L2 protein estimated using the LG+F+I+ρ4 substitution model. (C) Phylogeny of the Gammaherpesvirus estimated using the LG+I+ρ4 amino acid substitution model. Trees are midpoint rooted for clarity. Scale bars indicate number of amino acid substitutions per site. Viruses identified in this study are identified with red text and animal silhouettes indicate the host species. Branch support values >90% are indicated with a black dot at the node.

Two novel anelloviruses were detected, although only one of these – here named Antechinus anellovirus - contained the ORF1 sequence and hence was analysed further. Phylogenetically, Antechinus anellovirus was extremely divergent in the ORF1 protein from the current diversity of *Anelloviridae*, and was closely related to a group of anelloviruses isolated from seals, cats and a giant panda (Figure 6B). Antechinus anellovirus was detected in two libraries of spleen, co-occurring with Antechinus hepacivirus in one of these libraries.

**Figure 6.**
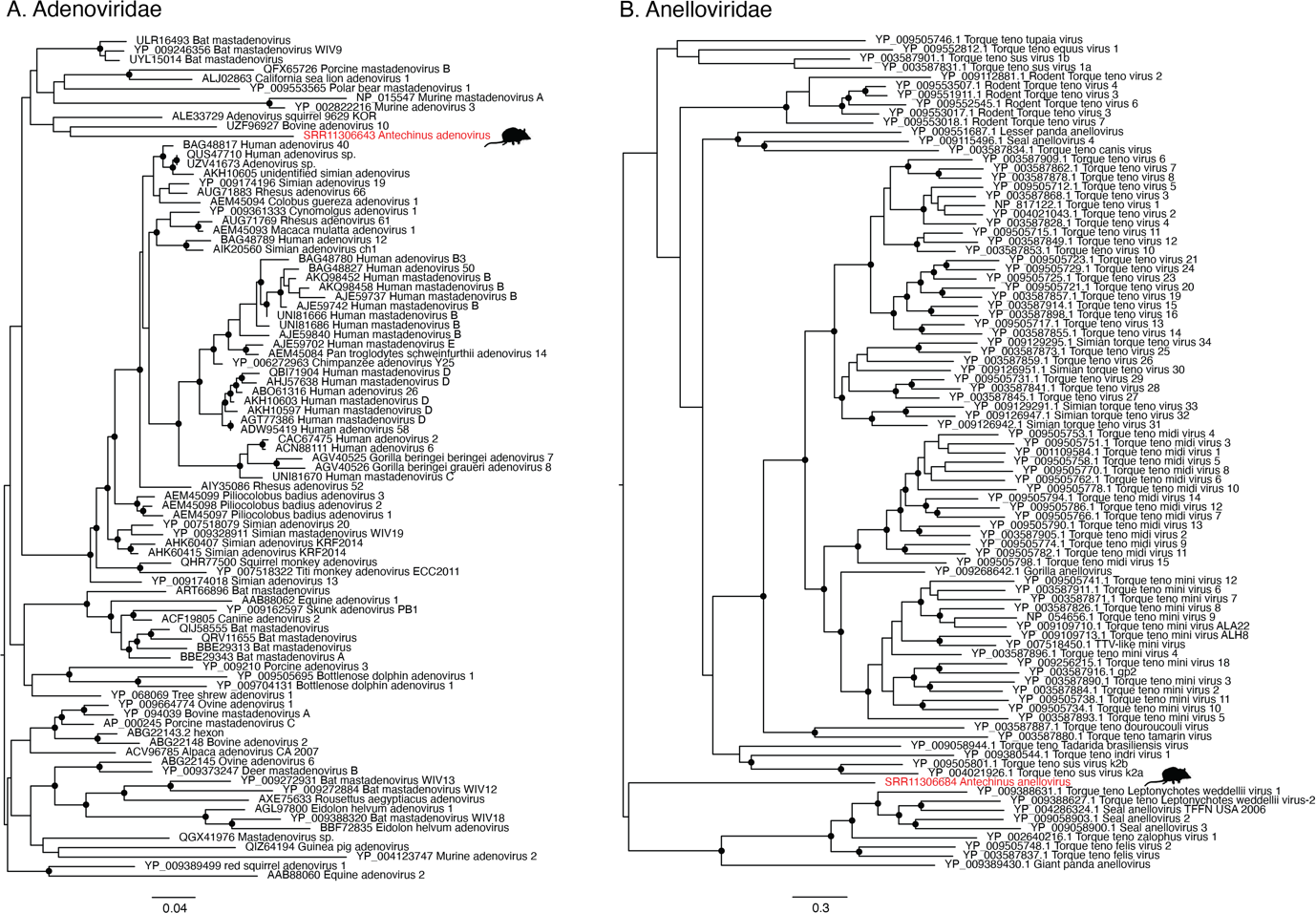
Phylogenies of *Adenoviridae* and *Anelloviridae*. (A) Phylogeny of the *Adenoviridae* hexon protein estimated using the LG+I+ρ4 substitution model. (B) Phylogeny of the *Anelloviridae* hexon protein estimated using the LG+F+ρ4 substitution model. Viruses identified in this study are identified with red text and animal silhouettes indicate the host species. Trees are midpoint rooted for clarity. Branch support values >90% are indicated with a black dot at the node. Scale bars indicate the number of amino acid substitutions per site Of interest, a library that contained the E2 protein of Tasmanian devil papillomavirus 5 (SRR13765816) also contained a contig showing sequence similarity to the large T antigen of a novel polyomavirus. A second library, SRR13765794, contained an identical polyomavirus large T antigen but no other virus sequences. No other polyomavirus genes were detected in these libraries. As this large T antigen sequence was related to, but distinct from Tasmanian devil polyomavirus 1 and 2 (Figure 5A), we named this novel virus Tasmanian devil polyomavirus 3.

Similarly, the adenovirus identified in this study was also found in the same samples as Antechinus hepacivirus: in the kidney tissue of two individuals, as well as in eight other libraries (seven individuals) where it was the only virus identified. Across libraries the virus sequences were 99.8% similar at the nucleotide level. The virus was mainly found in libraries of kidney tissue (8/10) as well as one library of spleen and one of prostate tissue. Antechinus adenovirus was most closely related to members of the genus *Mastadenovirus*, and based on a phylogeny of the hexon protein, grouped with samples isolated from bats, pigs and cows as well as a single squirrel sequence (Figure 6A).

The most diverse viral family identified were the *Papillomaviridae*. Specifically, we discovered five species of novel papillomavirus in six libraries of Tasmanian devil lip tissue from a study of Devil Facial Tumour Disease (DFTD) (Kozakiewicz et al. 2021). These viruses were named Tasmanian devil papillomavirus 3-7 to remain consistent with previously identified Tasmanian devil associated papillomaviruses (Chong et al. 2019). Partial gene transcripts were identified in all six libraries, although the gene transcripts present were not consistent across all libraries. The L2 gene (minor capsid protein) was found in all six libraries and thus used for comparison. Five of these were distinct papillomavirus L2 genes, with amino acid sequence similarity ranging from 36 – 59%; as the ICTV species demarcation for this genus is <70%, these were deemed to represent five distinct virus species. As the L1 (major capsid) protein was present in five libraries with distinct L2 transcripts, L1 and L2 were used for phylogenetic analysis. These sequences formed a distinct clade of marsupial papillomaviruses with a bettong papillomavirus and the bandicoot papillomavirus that is the sole member of the genus *Dyolambdapapillomavirus* (Figure 5B). The presence of papillomaviruses did not correlate with the DFTD status of the animal as both DFTD positive and negative samples contained papillomaviruses and there was no phylogenetic distinction based on DFTD status.

As the study for which these data were generated was designed to determine geographical patterns in DFTD expression profiles (Kozakiewicz et al. 2021), we assessed whether geographic location affected the phylogenetic clustering by manual inspection of the tree. No obvious correlation was observed. Two papillomavirus species were previously identified in Tasmanian devil faecal samples - Tasmanian devil-associated papillomavirus 1 and Tasmanian devil-associated papillomavirus 2 - although only the E1 protein was published for these viruses. Of the viruses discovered here, only a small fragment (317 nt) of the E1 region could be recovered for Tasmanian devil papillomavirus 6. This exhibited 48% aa similarity to Tasmanian devil-associated papillomavirus 1, 55% aa similarity to Tasmanian devil-associated papillomavirus 2, and 28% aa similarity Bettong penicillata papillomavirus 1.

## 4. Discussion

We identified 22 full or partial virus genomes from eight virus taxa from 446 SRA libraries of tissue samples from species within the Dasyuromorphia. Of these virus genomes, 15 were subjected to phylogenetic analysis. The remaining seven virus contigs were either viruses identified as contaminants (in four cases discussed below) or were too short for meaningful analysis; this was the case with two herpesviruses identified in a fat-tailed dunnart library and a Tasmanian devil library, as well as the ORF2 segment of a novel Anellovirus in a Tasmanian devil. Human papillomavirus sequences were identified in four antechinus libraries, three of which also contained contaminating human sequences. Although this is a risk when analysing SRA data, the use of taxonomic classification software such as CCMetagen (Marcelino et al. 2020) can be used to identify contaminating host sequences. We can also be confident that the viruses identified here are exogenous, rather than endogenous, as high-quality host genomes are available for three of the four Dasyurid species studied here, and a comprehensive analysis of endogenous viral elements in marsupials has previously been undertaken (Harding et al. 2021).

Until 2018, deltaviruses were believed to be exclusive to humans. However, this was disproven by the identification of delta-like viruses in birds and snakes (Hetzel et al. 2019; Wille et al. 2018). Since this time, additional deltaviruses have been identified through SRA screening (Bergner et al. 2021). Of note, we describe the first marsupial deltaviruses which also form what appears to be a marsupial specific clade. Specifically, Tasmanian devil deltavirus and Fat-tailed dunnart deltavirus cluster in a sister clade to a group of rodent associated deltaviruses also identified in a SRA screening study (Bergner et al. 2021). Further exploration of the marsupial virome is needed to determine if this putative marsupial cluster of deltaviruses holds true, such that these viruses have associated with marsupial hosts for their entire 160 million years history (Luo et al. 2011), or if there has been frequent cross-species transmission in other deltaviruses (Bergner et al. 2021). Two distinct marsupial carnivore hepaciviruses were also identified in this study. Hepaciviruses are associated with liver disease in humans, although in most animals their importance as agents of disease is unknown. Three marsupial hepaciviruses have previously been identified: Koala hepacivirus, Possum hepacivirus and Collins beach virus, all of which were identified metagenomically (Chang et al. 2019; Porter et al. 2020) (Harvey et al. 2019). Antechinus hepacivirus was most closely related to a suspected bandicoot associated hepacivirus, Collins beach virus, while Koala hepacivirus and possum hepacivirus cluster together. These two marsupial clades are paraphyletic, with Antechinus hepacivirus and Collins beach virus grouping with rodent hepaciviruses, while the possum and koala hepaciviruses cluster with a broader range of mammals. This may be a result of bandicoots and antechinus sharing a similar ecological niche – ground dwelling foragers (along with rodents) – thereby enabling cross-species transmission, while koalas and possums are mainly tree dwelling herbivores. The lack of a marsupial specific clade, as well as the grouping with rodent associated hepaciviruses, is also indicative of multiple introductions of hepaciviruses to marsupials, perhaps via rodent species, although this will again need to be resolved through more extensive sampling. Of note, Antechinus hepacivirus was identified in a relatively large proportion of individuals (5/13), which could suggest a high prevalence in the population and hence be of particular concern if populations are being used for translocation in population reseeding programs (Manning 2021). In addition, we identified a highly divergent hepacivirus in a numbat liver transcriptome. This virus shared only 29% amino acid sequence similarity to the closest blast hit and did not fall within the larger mammalian/marsupial clade suggesting the presence of a third, highly divergent lineage of marsupial hepaciviruses. Further study of marsupial associated hepaciviruses is needed to shed light on the emergence and evolution of these viruses in Australian marsupials.

A highly divergent chu-like virus was identified in a transcriptome of a DFTD cell line (SRR6380970) grown from primary tissues (Kosack et al. 2019). The *Chuviridae* and Chu-like viruses belong to the order *Jingchuvirales*, which was until recently believed to be an invertebrate specific order (Di Paola et al. 2022). The discovery of a chuvirus in the brain of a snake with neurological disease (Argenta et al. 2020) and in a meta-transcriptomic study of reptiles (Shi et al. 2018) led to the identification of a fish-reptile associated genus (*Piscichuvirus*) within the *Chuviridae*. More recently, three novel *Piscichuvirus* species were identified and associated with encephalitis in turtles (Laovechprasit et al. 2023), suggesting that this genus could be associated with neurological disease. Tasmanian devil chu-like virus is the sister-group to the *Chuviridae* within the Jingchuvirales and is highly divergent from the characterised species. Interestingly, it has been suggested that DFTD is of Schwann cell origin (Murchison et al. 2010), a cell type associated with the peripheral nervous system (PNS), such that the novel chu-like virus identified here could similarly be associated with the nervous system. However, as this virus was identified from SRA data and from a library of cultured cells, we cannot be certain of the true host of this virus.

Five families of DNA viruses were identified in these samples. Papilloma-, polyoma- and herpesviruses are often associated with disease and, in some cases cancer, while anellovirus and adenovirus have less definitive links to disease. We identified a novel anellovirus in antechinus spleen transcriptomes, which co-occurred with Antechinus hepacivirus in one library. These viruses are ubiquitous and their disease association is still debated, although it has been suggested that they may play a role as co-infecting viruses (Webb et al. 2020). Similarly, Antechinus adenovirus was identified in kidney, prostate and spleen tissue transcriptomes and co-occurred with hepacivirus in two of ten libraries. This virus clustered with a group of mastadenoviruses identified in mice, cows, pigs, bats and a polar bear. Mastadenoviruses are in some cases associated with disease including encephalitis, respiratory disease (Chen and Tian 2018), and gastrointestinal symptoms (Dayaram et al. 2018). In this case, no disease was reported in the individuals sequenced, but this is difficult to assess in SRA screening studies.

Two distinct species of gammaherpesvirus were identified in antechinus transcriptomes of stomach and prostate tissue, along with partial herpesvirus sequences in Tasmanian devil peripheral blood mononuclear cells and fat-tailed dunnart mitogen-stimulated splenocytes. Herpesviruses are relatively well characterised in marsupials, with Dasyurid herpesvirus 1 identified in *Antechinus* in 2014 (Amery-Gale et al. 2014) and subsequent studies revealing two Tasmanian devil associated herpesviruses (Dasyurid herpesvirus 2, Dasyurid herpesvirus 3) (Chong et al. 2019; Stalder et al. 2015). However, the species demarcation of viruses within the family now termed *Orthoherpesviridae* is unclear, with no defined genetic distance or biological features used for classification (Gatherer et al. 2021). Given that very few genomic sequences are available for the related viruses, we deemed 92% nucleotide similarity across the single read of the DNA-DNA polymerase (the only gene available for Dasyurid herpesvirus 1) as sufficient genetic distance to represent a novel species. DNA sequencing would likely lead to the recovery of the full genome, including gene sequences necessary to determine the relatedness of these species to each other and to other dasyurid herpesviruses, but this is beyond the scope of this study.

We identified five distinct but related species of papillomavirus in Tasmanian devil lip tissue transcriptomes within the libraries of a single SRA BioProject. The DFTD status of the sampled individual did not appear to correlate with the presence of papillomavirus, nor did DFTD status correlate with any clustering in the phylogeny of these species. Two Tasmanian devil associated papillomaviruses were previously identified in a faecal meta-transcriptome, but as these animals are carnivores it was impossible to determine their true host (Chong et al. 2019). Phylogenetic analysis revealed that all these sequences were most closely related to each other and more broadly to Bettong penicillata papillomavirus 1, isolated from papillomatous lesions on a brush-tailed bettong in Australia (Bennett et al. 2010), and more distantly with Bandicoot papillomatosis carcinomatosis virus type 1 and 2. Bandicoot papillomatosis carcinomatosis virus uniquely exhibits genomic features of both papillomavirus and polyomavirus proteins with the L1 and L2 proteins of a papillomavirus and the large and small T antigen proteins of a polyomavirus (Woolford et al. 2007).

Interestingly, the large T antigen protein sequence of a novel polyomavirus was identified in one library in this study which also contained the E2 protein of Tasmanian devil papillomavirus 5, although this is not consistent with the genomic structure of the previously identified hybrid virus. Phylogenetically, the large T antigen sequence of Tasmanian devil polyomavirus 3 was most closely related to the large T antigen of the hybrid Bandicoot papillomatosis carcinomatosis virus, with Tasmanian devil-associated polyomavirus 1 and Tasmanian devil-associated polyoma-like virus 2 more divergent. Interestingly, Tasmanian devil-associated polyomavirus 1 and Tasmanian devil-associated polyoma-like virus 2 were identified in the same faecal meta-transcriptome study as Tasmanian devil-associated papillomaviruses 1 and 2. DNA sequencing will be required to confirm the genomic structure and phylogenetic relationships of these viruses.

Although our study identified a relatively high diversity of DNA viruses within transcriptomic data, the lack of complete genomes limits the value of these sequences in phylogenetic comparison and genomic characterisation of these species. In contrast, there was a relatively limited diversity of RNA viruses in these data which may be a product of the sequencing methodology used. For example, poly-A selection during library preparation will remove many virus sequences (Visser et al. 2016). However, despite being prepared with a poly-A selection, we identified a high diversity of viruses within the yellow-footed *Antechinus* libraries, with 10/13 sequenced individuals being associated with at least one virus. Antechinus hepacivirus was particularly prevalent and as these species are being proposed for species reintroduction programs, it is of particular importance that the viromes of these individuals are investigated to prevent the spread of these viruses to naive populations.

In light of the diversity of marsupial carnivore viruses identified in this small study of transcriptomic data not generated for the purposes of virus discovery, it is evident that considerable viral diversity exists in these species and that further study is required to understand the viromes and viral evolutionary history of these unique species. Given the threat that these species face as a result of human activity, it is imperative that we understand the risk that viral disease poses to what are often small, isolated populations.

## Acknowledgements

We acknowledge the University of Sydney’s high-performance computing cluster, Artemis, for providing computing. This work would not be possible without the sequencing data that have been generously shared by the research community, and we are grateful to the NCBI SRA team for their management of this resource. This work was funded by an National Health and Medical Research Council Investigator Grant to ECH (GNT201719).

## Data availability

All viral genomes and corresponding sequences assembled in this study have been deposited in GenBank under BioProject XXXXXXX, and with accession numbers XXX-XXX. The alignments generated in this study for phylogenetic analysis are available at https://github.com/erinhunter4/dasyurid_virome

