## Supplementary figures and images for "Divergent hepaciviruses, chuvirus and deltaviruses in Australian marsupial carnivores (Dasyurids) identified through transcriptome mining"

### Supp. Fig. 1

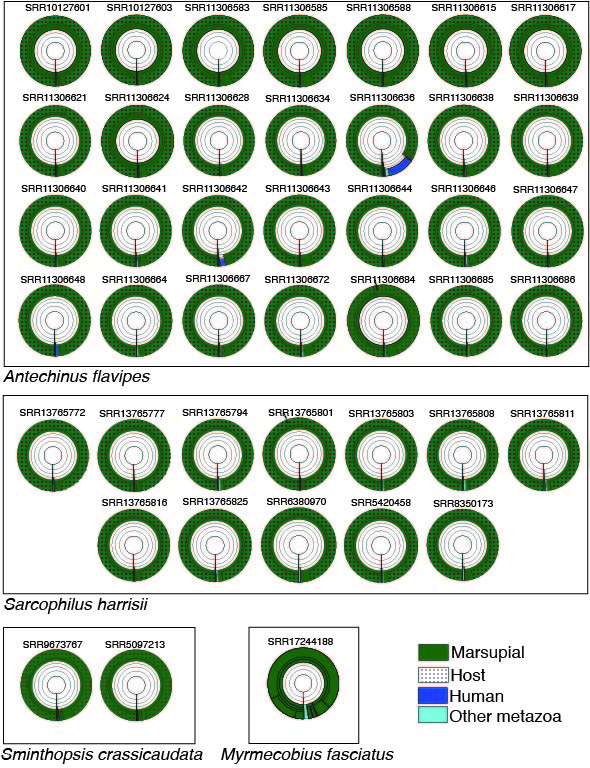
